# Modelling biological and resource fluxes in fluvial meta-ecosystems

**DOI:** 10.1101/2024.01.12.575367

**Authors:** Lauren Talluto, Rubén del Campo, Edurne Estévez, Thomas Fuß, Lukas Thuile Bistarelli, Jan Martini, Gabriel A. Singer

**Affiliations:** Biodiversity, Ecology, and Conservation Group, International Institute for Applied Systems Analysis, Laxenburg, Austria; Department of Ecology & Hydrology, University of Murcia; Department of Ecology, University of Innsbruck, Innsbruck, Austria; Faculty of Science and Technology, University of the Basque Country (UPV/EHU), Spain; gutwasser GmbH, Geerenweg 2, 8048, Zürich, Switzerland

**Keywords:** fluvial meta-ecosystems, biodiversity, ecosystem functioning, nutrients, river networks

## Abstract

Meta-ecosystem theory predicts that cross-ecosystem flows of energy, nutrients, and organisms have important implications for local community assembly and ecosystem functioning. Developments in the theory also have the potential to enhance our understanding of biodiversity-ecosystem functioning relationships. Meta-ecosystem theory is particularly well-suited to the study of rivers, because water flow forces strong spatial interrelationships among connected ecosystems. However, models that address flows of both resources and organisms and explicitly link both are lacking. We present a model and associated R-package for cross-ecosystem flows of both resources and organisms that can be used to predict their distribution in river networks, as well as meta-ecosystem functioning. The model incorporates feedbacks between these two crucial components—resource concentrations represent niche dimensions for organisms, modifying the colonisation and extinction dynamics and different locations in the network, and organisms also consume resources, thereby modifying the concentrations that are transported downstream. To illustrate the capabilities of the model, we present an in silico experiment and analysis, as well as providing sample code.

## 1 Introduction

Understanding the effects of biodiversity on ecosystem functioning (EF) has long been a central goal of ecological research (Bannar-Martin et al. 2018). In general, biodiversity is an important driver of EF (Cardinale et al. 2012; Hooper et al. 2012) and can be as important as abiotic drivers (e.g., climate Duffy et al. 2017), leading many researchers to hypothesise that ecosystems with more species would exhibit higher levels of EF (Margalef 1963). However, recent experiments and models have revealed complex effects of diversity on EF (i.e., positive, negative and no effects; see Pennekamp et al. 2018), pointing to non-linear and context dependent biodiversity-EF relationships (Thébault and Loreau 2006; Little and Altermatt 2018). This further suggests additional important drivers of EF beyond the diversity of organisms. For example, the rates of acquisition, use and allocation of resources are key EFs that can depend on species composition (and corresponding functional traits) due to the effects of morphology, behaviour, and physiology (Hooper et al. 2005; Díaz et al. 2013; Gagic et al. 2015). Moreover, different resources contain different essential nutrients and carbon compounds that determine lability and palatability; thus resource properties can also strongly determine consumption and processing rates (Cornwell et al. 2008).

River networks are characterised by high biodiversity, variability in spatial structure, and large fluxes of energy, nutrients, and organisms among ecosystems. This is in large part due to the overwhelming imprint that the flow of water has on the spatial structure of fluvial networks. In particular, the relative isolation of less-connected upstream portions of a fluvial network contrasts strongly with downstream areas where water flow brings organisms and nutrients from many upstream regions. Moreover, because water flow is temporally variable, the degree of connectivity is similarly dynamic. Finally, rivers are embedded in a terrestrial environment, implying that the physical and geographical structure of river networks promotes the exchange of resources and organisms both between aquatic and the surrounding terrestrial ecosystems, as well as among different locations in the river (Cid et al. 2022; Battin et al. 2008). These processes have profound implications for community assembly and EF. The spatial structure of a river network, and the degree to which this structure differentially influences the movement of organisms and materials, may significantly impact EF at the scale of the entire network (Talluto et al. 2024). Recent developments in the theory of meta-ecosystems provide a solid conceptual framework for describing how both organisms and material interact in such a spatial context, and how this interaction can influence the spatial distribution of EF (Loreau et al. 2003; Gravel et al. 2010; Harvey et al. 2023).

Despite developments in theory, field studies of meta-ecosystems are challenging due to the complexities of the processes, the disconnect between theoretical processes and what is feasible to measure in the field, and the multiple spatial scales generally involved in most meta-ecosystems (Gounand et al. 2018). In order to better integrate the theory of meta-ecosystems with actual field studies, we need both (i) highly resolved spatio-temporal data on resources and organisms within fluvial meta-ecosystems (FMEs), and (ii) a new generation of models operating at the meta-ecosystem level (Talluto et al. 2024). These models should link community composition with resources and EF. The availability of datasets is rapidly increasing thanks to technological developments that allow for greatly improved resolution in studying both community composition (e.g., through eDNA and metabarcoding) as well as resource composition diversity (e.g., size-exclusion chromatography, mass spectrometry and infrared spectroscopy) (Sleighter and Hatcher 2008; Huber et al. 2011; Tremblay et al. 2011; Baird and Hajibabaei 2012). These datasets are unprecedented in both breadth—covering a great spatial extent, such as entire river networks—and depth—capturing much more biological and chemical diversity at a single location than has ever been possible (Altermatt et al. 2020). However, we still lack models that can provide the context for these datasets at the appropriate spatio-temporal scales (Gounand et al. 2018; but see Harvey et al. 2023).

Developing fluvial meta-ecosystem models presents important challenges due to the unique properties and structure of river networks. For example, the spatial configuration of river networks imposes unique and strong controls on both organisms and resource fluxes. However, the distribution of these ecosystem components is driven by different mechanisms. Resources highly depend on structure of the surrounding terrestrial ecosystem (i.e., land cover shapes the nature of the resources that are transferred to the river corridor) and are passively transported by flow (Roebuck et al. 2020). In contrast, the distribution of organisms is driven by the interplay between (i) stochastic processes (e.g., extinctions); (ii) dispersal (including both flow-mediated downstream dispersal but also active dispersal and transport by other organisms); (iii) local environmental factors, including resource properties, that filter species from the regional species pool; and (iv) biotic interactions (Leibold et al. 2004; Urban 2004; Heino et al. 2015).

Here, we develop a meta-ecosystem model that explicitly couples the distribution and fluxes of both resources and organisms in FMEs. We construct this model by joining two commonly-used model types in fluvial ecosystem and community ecology: transport-reaction models (for resources), and metacommunity models (for organisms). By linking these two models, we allow for feedbacks between the organisms and the resources they use. These feedbacks are essential to enable a more mechanistic understanding of the connections between the processes underlying community assembly, resulting biodiversity patterns, and EF. Our model structure facilitates developing and testing *in-silico* hypotheses regarding FME properties. These properties can include, for example, interactions of biological communities with resources in the context of river spatial structure, and both local- and regional-scales, and the consequences of these iterations on patterns in biodiversity and EF, The model also allows for manipulations of river connectivity, flow, resource and organism traits, and the terrestrial matrix. We provide an implementation of the model via an R-package as well as some worked examples with sample code and data.

## 2 Materials & Methods

### 2.1 Model Description

Our conceptual starting point for modelling the biological community is metapopulation theory (Levins 1969). The classic metapopulation model tracks the number of occupied patches *p* in a landscape composed of *h* available patches as a function of the rates of colonisation and extinction of local populations within those patches:

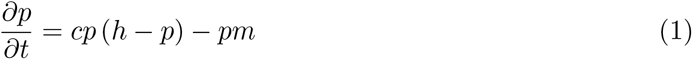

In this model, extinction occurs within occupied patches according to the extinction rate *m*, resulting in the total negative occupancy flux of *pm*. Similarly, unoccupied patches are colonised according to the colonisation rate *c*. The additional *p* in the colonisation term takes dispersal into account; as more patches are occupied in the metapopulation, unoccupied patches are colonised more quickly due to increased dispersal from the occupied patches.

Hunt and Bonsall (2009) extended this model to multi-species communities by adding competition to the extinction term:

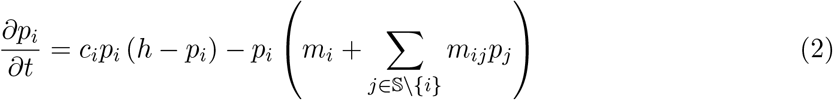

where the subscript *i* indicates a focal species, the subscript *j* a competitor, and 𝕊 \ {*i*} is the set of species in the metacommunity excluding the focal species *i*. Here, the extinction rate is now broken into two terms. The first a species-specific intrinsic extinction rate *m*_*i*_, representing, e.g., stochastic extinctions that are unrelated to other species. The *m*_*ij*_ term is the effect of competition between *i* and *j* on the extinction rate of *i* (multiplied by *p*_*j*_ because competition only occurs when species *j* is also present). For future clarity of notation, we assume all parameters are specific to a target species *i* and thus omit the subscript.

The model in equation 2 describes regional dynamics; the rates of dispersal, competition, colonisation and extinction are constant across all locations in the metacommunity. However, these rates certainly vary in space in real metacommunities, particularly in rivers where habitat conditions and river flow follow strong gradients (e.g., discharge generally grows moving from upstream to downstream). We first consider how river flow interacts with dispersal in local communities (denoted by the subscript *k*). In rivers, hydrological connections among habitat patches facilitate passive dispersal with flowing water. In addition, many organisms are capable of active dispersal (i.e., any dispersal not facilitated by the flow of water in the river channel), either overland or upstream along the river course, and even organisms that cannot move themselves can be carried upstream by various processes (e.g., transport by other organisms, wind, or humans; Kristiansen 1996). Because these two dispersal modes are likely to occur at quite different rates, and because the rates will likely vary among different types of organisms, we break the dispersal portion of the colonisation term here into active (*α*) and passive (*β*) components, with the passive component additionally weighted by the input discharge *Q*_*k*_. Finally, local dispersal is weighted by the occupancy of reaches within dispersal range (denoted with *k*^t^).

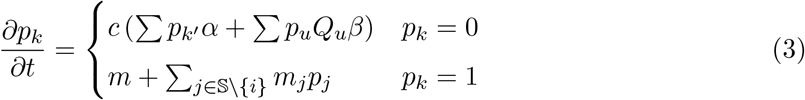

It should also be noted that, since this equation is local, only one of the colonisation or extinction flux will apply at any given time, depending on whether the site is occupied.

An additional important factor is variation in the local environment, which could contribute both to colonisation and extinction fluxes. The Levins model (eqn. 1) has been extended by more recent theoretical (Holt and Keitt 2000; Holt et al. 2005) and empirical (Talluto et al. 2017) work by fitting the colonisation and extinction rates as functions of local climatic conditions. The result is a dynamic range model with long-term occupancy driven by the balance of local colonisation and extinction (Talluto et al. 2017). We combine this approach with the multi-species Hunt model (eqn. 2) by redefining the *c* and *m* terms to be functions of the quality *q*_*k*_ of the focal patch:

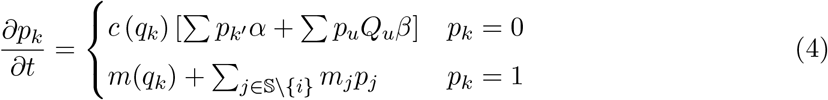

The shape of these two functions is flexible; for example, Talluto et al. (2017) defined them as quadratic functions of local climate conditions. Our more general approach here can incorporate local climatic and/or habitat conditions, and dynamic resource concentrations into the *q* term. We explore these possibilities further in the case studies. For now, we consider the case where *q*_*k*_ = [*R*]_*k*_, the concentration of an essential resource. We can use a simple transport-reaction model (Soetaert and Herman 2009) to describe the fluctuations of this resource concentration in time and space (since all terms relate to a focal patch, the subscript *k* has been omitted for clarity).

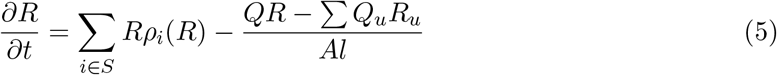

The first term on the right hand side models reaction; we consider here only reaction due to biological activity, and postulate that the net reaction in a patch is the sum of a set of resource use functions, *ρ*_*i*_(*R*), of all species 𝕊 in the local community. Each *ρ*_*i*_ function describes the impact of a species on the resource in units of *s*^−1^. The forms of these functions will depend on which resources are being modelled. The second term gives loss due to advective transport, where *Q* is the discharge of the focal patch (in volumetric units per unit time; we use m^3^ s^−1^ throughout). Each patch receives inputs both in the form of lateral flow (e.g., overland flow, groundwater, etc.) as well as flow from upstream reaches. Thus, the total input resource mass for the focal patch will be the sum of input masses, which are the product of the input concentrations (*R*_*u*_) and discharges (*Q*_*u*_). Finally, the masses are converted back to a concentration using the cross-sectional area (*A*; m^2^), and length (*l*, m).

### 2.2 Implementation

We provide an implementation of the model in an R-package **flume** (FLUvial Meta-Ecosystem model) (Talluto 2023), and demonstrate some capabilities via case studies and sample code. In this section we describe some of the implementation details that are flexible in the formulation of the model above, but for which we have made specific choices in software. A full list of model parameters and default values from the package is available in Table 1.

**Table 1:** Model parameters with defaults and values used for case studies.

| Symbol | Description | Default Value | Case Study 1 | Case Study 2 |
| --- | --- | --- | --- | --- |
| $c_\mu$ | Niche location | $0 - 1^*$ | $0.1 - 0.9$ | varies $^\ddagger$ |
| $c_\sigma$ | Niche breadth | $c_{\mu,2} - c_{\mu,1}^\dagger$ | 0.1 | varies $^\ddagger$ |
| $c_\gamma$ | Colonisation scale | $5 \times 10^{-6}$ | $6 \times 10^{-6}$ | varies $^\ddagger$ |
| $m_\gamma$ | Extinction scale | $2 \times 10^{-7}$ | $1.25 \times 10^{-7}$ | varies $^\ddagger$ |
| $\alpha$ | Active dispersal rate | 0.1 | $0.05 - 2.0$ | 0.5 |
| $\beta$ | Passive dispersal rate | 0.1 | 0.1 | 0.1 |
| $o_\gamma$ | Competition scale | 0.001 | 0.001, 0.1 | $1 \times 10^{-4}$ |
| $r$ | Resource use scale | $5 \times 10^{-4}$ | $5 \times 10^{-4}$ | $1 \times 10^{-3}$ N, $5 \times 10^{-4}$ P |

\* Uniformly distributed across the range, depending on the number of species.
$^\dagger$ i.e., the distance between niche locations of adjacent species.
$^\ddagger$ Parameters determined empirically, see §3.2

#### 2.2.1 Patch quality

Patch quality is defined by two functions describing the impact of quality on colonisation and extinction (eqn. 4). We modelled the colonisation niche (i.e., the shape of the *c* function) as a Gaussian curve (Austin 1999). Thus, for a single resource, the niche shape can be defined by three parameters: location (*c*_*µ*_), breadth (*c*_*σ*_), and scale (*c*_*τ*_ ):

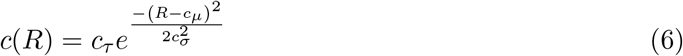

With multiple niche dimensions, the niche shape is similarly described by a multivariate Gaussian. Extinctions in the model can be constant with respect to the environment, controlled only by a scale parameter (*m*_*γ*_), or can follow an inverse Gaussian with a location, breadth, and scale.

#### 2.2.2 Competition

Our metacommunity model includes a competition term (∑*m*_*i*_*jp*_*j*_), allowing for totally independent pairwise (and potentially asymmetrical) competition coefficients among species. In practice, pairwise competition strength is extremely difficult to estimate. Thus, we make a simplifying assumption that the strength of competition will be proportional to the degree of overlap in two species’ fundamental niches (defined as the difference between colonisation and extinction, excluding the competition term). We further add a scale parameter *o*_*γ*_ (which is constant by default but can vary by species pair). In flume, we compute this as the area of overlap in the two species’ niches, represented mathematically as the integral of a piecewise function defined by the lesser of the two species’ niches when both niches are positive (i.e., within the fundamental niche of the species), and zero otherwise:

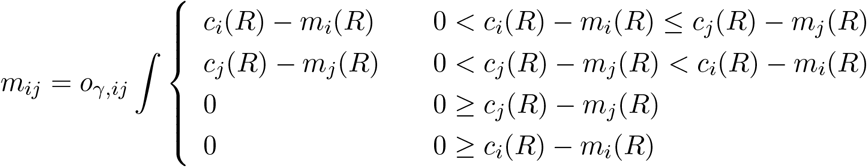

#### 2.2.3 Resource use functions

In principle, we expect that a species’ total resource consumption at a patch will be correlated with its abundance (or biomass) (Díaz et al. 2003; Winfree et al. 2015). Although we do not track abundance in our model, we can make the simplifying assumption that abundances will be higher when species are closer to their niche optima (Holt et al. 1997; but see McGill 2012). Thus, the consumption *ρ*_*i*_(*R*_*k*_) by species *i* at site *k* (eqn 5) will be proportional to the niche height at that site *c*_*i,k*_ − *m*_*i,k*_ (eqn. 4). We can further allow that species vary in their ability to use resources, scaling the resource use function by a species-specific consumption constant *r*_*i*_:

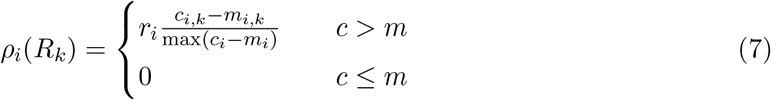

Here we scale the niche height by the maximum niche height to allow us to interpret this term as the fraction of a species’ maximum consumption rate, and to interpret *r*_*i*_ as the maximum rate. Finally, we assume that when a species is outside its equilibrium habitat (i.e., when *m* > *c*; Talluto et al. 2017), resource consumption is negligible and can be ignored.

#### 2.2.4 Stochastic simulations

For analysis, we discretised the model in space and time. We thus consider a series of habitat patches representing stream reaches. Each reach is characterised by two fundamental state variables: the resource vector **R**, giving the concentration of each resource, and the community vector **C**, which gives the absence (denoted by zeroes) or presence (denoted by ones) of all possible species in the community. For any time interval Δ*t* we can then compute the probability of observing colonisations and extinctions for a species *i* (eqn. **??**):

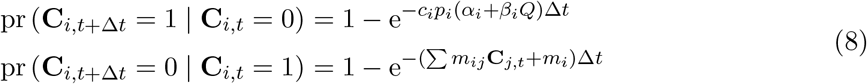

Here, the prevalence term *p*_*i*_ indicates local prevalence, i.e., the number of occupied reaches within dispersal range. We consider only the case of nearest neighbour dispersal, so *p*_*i*_ is the sum of the occupancy states of reaches immediately up- and downstream of the focal reach; longer distance dispersal can be incorporated via the regional species pool (see §**??**).

Changing resource concentration can be computed for each reach using a variety of numerical integration techniques. The flume package uses the lsoda algorithm implemented in the DeSolve R-package (Soetaert et al. 2010).

### 2.3 Boundary Conditions

Boundary conditions in the model are used to represent external input into the river network (e.g., input of resources and organisms from the terrestrial environment, or overland flow of water and groundwater contributing to discharge). Rivers are not closed systems, and because of the downstream flow of water, all material would be rapidly flushed downstream without user-provided boundary conditions. In the model, these inputs are provided by a virtual reach with a constant user-defined resource composition and community composition. Thus, model terms that incorporate water flow from upstream (e.g., resource transport and passive dispersal) will incorporate inputs both from upstream in the river as well as from outside the river network. Lateral discharge from these reaches is not user supplied, rather it is computed as the growth in discharge from one reach to the next moving downstream, according to the growth in catchment area and defined by hydraulic scaling relationships (Burgers et al. 2014). By default, flume uses boundary conditions as initial conditions as well, but this can be overridden. The actual resource concentration and community composition of the virtual reaches can vary depending on modelling needs; for example, community composition could be uniform and contain all possible species to represent a classical “regional species pool” from metacommunity modelling, or resource concentrations could vary among headwaters or from upstream to downstream to represent land use gradients.

### 2.4 Temporal resolution

The time scale of the model determines how frequently colonisation and extinction steps are performed; community state is constant within time steps. The temporal resolution is set via a parameter passed at model initialisation, with units determined by the units of discharge. The default value is 86400, which, assuming discharge is in units of *m*^3^*s*^−1^, corresponds to 24 hours. Thus, using default parameters, community composition can change once per day, while resources are integrated continuously throughout the day. Longer time steps can be set to simulate slower community dynamics and improve computation time, with the potential downside that this allows resource concentrations to change more while community composition is constant.

### 2.5 Model output

After a simulation run, the model returns an object containing all model parameters and complete state (i.e., the model is resumable). The object also contains a complete history of all state variables, consisting of reach by time by resource and reach by time by species arrays. Resource fluxes (transport and consumption) are also returned for every reach and time step and are the primary metrics of EF returned by the model. The package also includes a variety of convenience functions for plotting and for computing aggregate metrics for both community and resource state.

### 2.6 Sample river network

In order to demonstrate some features of the package and explore the behaviour of the model, we created a representation of the Kamp river in Austria to use as the spatial setting for simulations. This river network is included with the package, along with other sample networks, and a vignette is included to help users generate a network from a digital elevation model.

## 3 Case studies

### 3.1 Case Study 1: Effects of dispersal and competition on biodiversity-EF relationships

For local communities, we expect that a strong match between environmental/resource conditions and the niches of occurring species should result in efficient use of resources and thus high local EF. It follows that, in order to achieve such a strong match, species that have appropriate niches must both exist and have the ability to reach local sites with the right conditions. A large regional species pool (i.e., high *γ* diversity) makes it more likely that species with fitting niches are available; thus at the network scale we expect *γ* diversity to correlate with the strength of biodiversity-EF relationships (i.e., networks with high *γ* diversity will have, on average, local communities with stronger biodiversity-EF relationships). However, *γ* diversity can only increase functioning if species are able to get to appropriate habitats. Thus, dispersal may be an essential mediator of biodiversity-EF relationships. If species are dispersal limited, then many niches may be vacant, resulting in local communities that cannot efficiently process resources and thus have low EF. On the other hand, high dispersal can lead to communities structured primarily by mass effects, resulting in communities with many redundant species or with species that are not well-adapted to local conditions (Mouquet and Loreau 2003; Declerck et al. 2013; Horváth et al. 2016). In other words, dispersal being either too low or too high can lead to a mismatch between environment and species and may reduce the strength of biodiversity-EF relationships.

We developed scenarios to see whether flume can reproduce the expectation that local-scale biodiversity-EF relationships are stronger at intermediate dispersal. We included *γ* diversity and the strength of competition as additional variables, to determine if they mediate any effect of dispersal on biodiversity-EF relationships. Using the Kamp river network (Fig. 2) as the spatial template, we started with a template metacommunity consisting of 50 species. All species had identical niche parameters (Table 1), with the exception of the niche location, which was assigned following a uniform sequence on the interval (0.1, 0.9). We scaled resource concentrations such that they were approximately between 0 and 1, for simplicity. From this template, we generated scenarios consisting of either a high-(40 species) or low-(20 species) diversity metacommunity. These scenarios were generated by drawing species randomly from the template metacommunity; for each diversity scenario, we generated a total of ten metacommunities this way, each with different randomly drawn species (Fig. 3).

**Figure 1:**
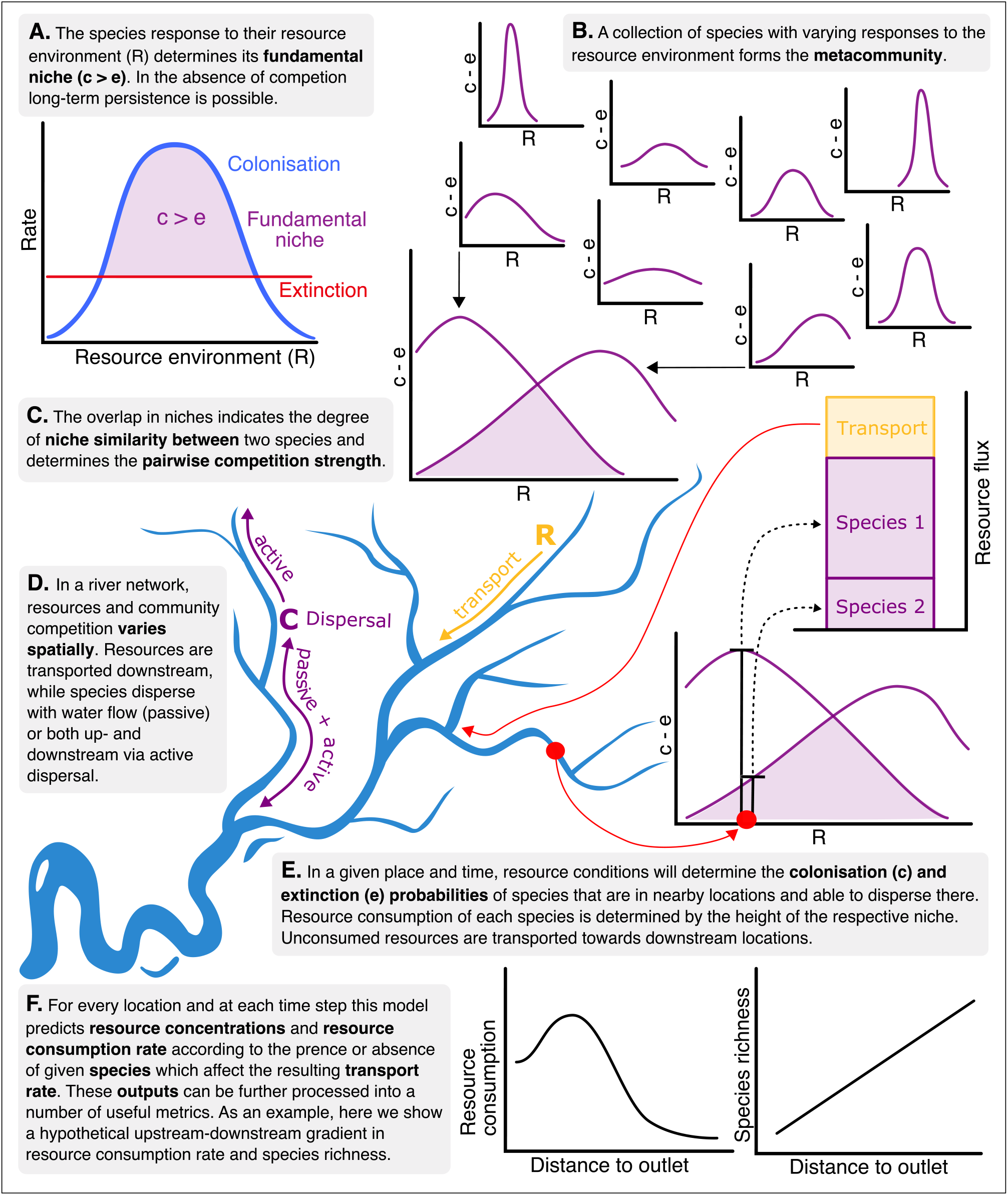
Conceptual diagram illustrating key model flows and outputs.

**Figure 2:**
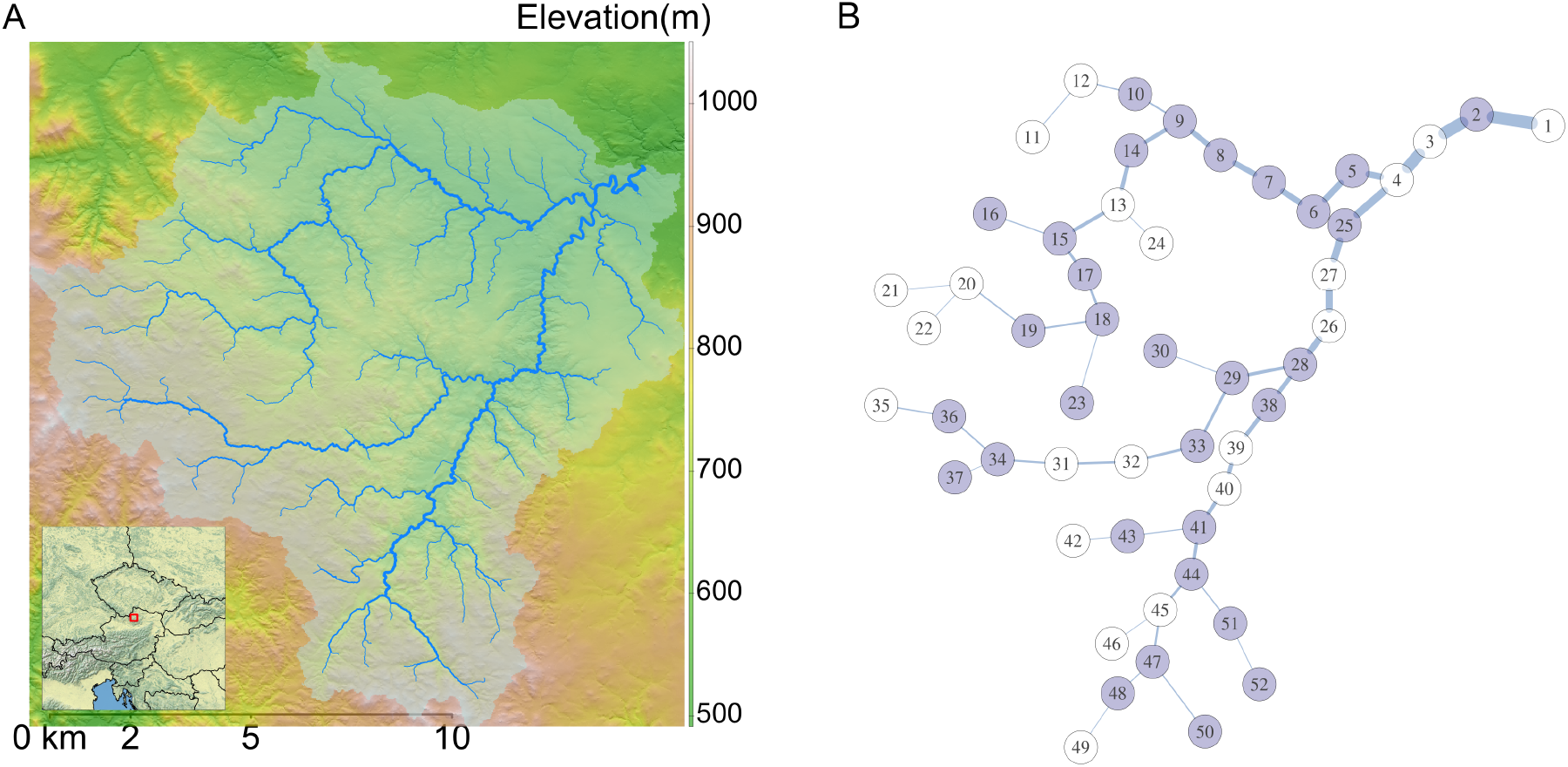
(A) Delineated Kamp river network, with the drainage area shown shaded. Line weights are proportional to Strahler stream order. Location of the river network within central Europe is shown in the inset. (B) Schematic of the Kamp river network after conversion for use with flume. Line weights are proportional to discharge, which ranges from 0.05–0.88 *m*^3^*s*^−1^. The network has been segmented into approximately 2-km reaches (which are assumed to be exactly equal length for modelling purposes); each node in the schematic is a reach midpoint. Reaches containing sampling sites (used in Case Study 2) are highlighted in purple.

**Figure 3:**
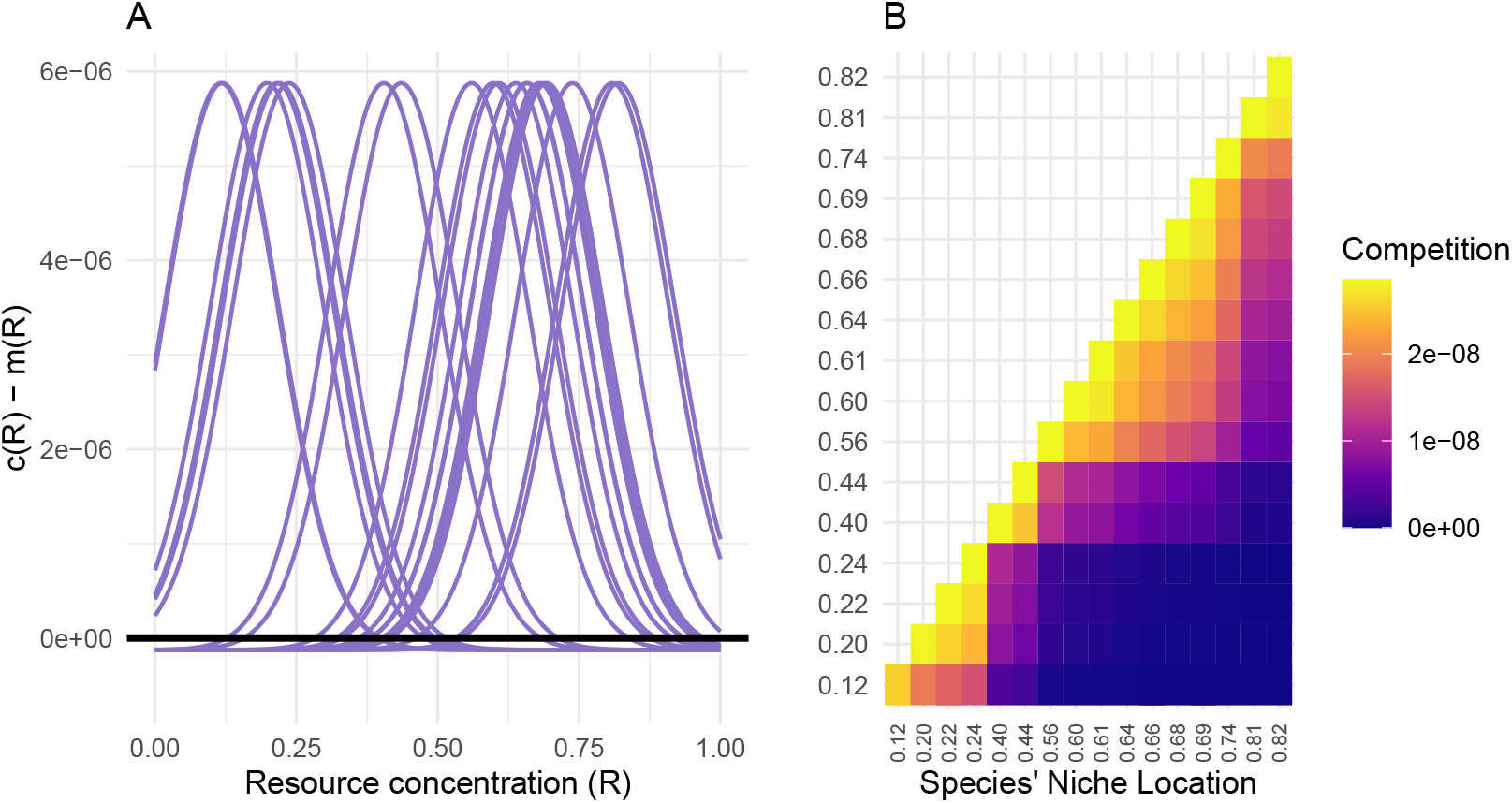
Visualisation of (A) niches and (B) strength of competition for a sample metacommunity of 20 species. In (A), the vertical axis is the difference between the colonisation term *c*(*R*) and the extinction term *m*(*R*). The height of this curve represents the net rate of change of occupied patches for given resource conditions for each species. Values greater than zero indicate conditions under which the species is able to persist in the long-term and represent the species’ fundamental niches. In (B), species are ordered following their niche location (i.e., the location on the resource axis of the optimum in A). Colours indicate pairwise competition strength, which are proportional to the degree of niche overlap.

For each of the ten randomly generated metacommunities, we ran simulations while varying dispersal ability and the strength of competition. Competition strength was varied by adjusting the competition scale parameter. In this way, local competition was always dependent on the number and niche parameters of species present, but globally competition was scaled up or down according to the scenario. We ran a high-competition scenario (*o*_*γ*_ = 0.1) and a low-competition scenario (*o*_*γ*_ = 0.001). We tested eight dispersal scenarios using the active dispersal parameter (*α*), setting *α* to the values (0.05, 0.1, 0.3, 0.5, 0.8, 1.2, 1.6, 2.0). The *α* = 0.05 was included as a “nearly zero” dispersal scenario; we used a small value instead of zero to avoid species being permanently lost from upstream reaches due to no upstream dispersal (and no immigration from outside the river network). The passive dispersal parameter was held at a constant low level (*β* = 0.1). All combinations of competition, *γ*-diversity, species lists, and dispersal ability resulted in 320 total scenarios. Additionally, to capture variability in the stochastic model, we replicated each individual scenario 8 times. All scenarios were run at a temporal resolution of 1 day, and for a total of 120 time steps. Boundary resource concentrations (i.e., the concentration of input from the surrounding land) were set to vary between 0.1 and 0.9 and were set to simulate a resource gradient due to, e.g., land use differences across the river network (Fig S1). Initial resource concentrations were set to match the concentration of input from the boundary conditions; we found in initial testing that, in the absence of fluctuations due to community changes, initial conditions had very little impact on model state since the river network quickly reaches an equilibrium resource state. Initial species locations were determined randomly with a per-species target prevalence of 20% of reaches occupied. For reporting results, we use a temporal average from the final 30 time steps. Taking the last time steps allows the model time to “forget” initial conditions, while averaging over time reduces some noise simply due to random fluctuations in colonisation/extinction of species. We use the total resource consumption rate (i.e., in units of *mass* × *volume*^−1^ × *time*^−1^) as the measure of EF, and species richness as the measure of biodiversity.

The extent to which dispersal reduced the efficiency of resource processing depended both on *γ* diversity and competition strength. The strength of the biodiversity-EF relationship also depended both on *γ* diversity and competition strength. When competition was weak, we observed a weak negative relationship between the biodiversity-EF slope and the dispersal ability, regardless of community diversity (Fig. 4). For both of these scenarios, there was no hump shape; i.e., the theoretical prediction that dispersal facilitates strong biodiversity-EF relationships up until mass effects result in poorly adapted communities was not supported when competition was weak. In contrast, with strong competition, we observed a pronounced hump in the relationship between the biodiversity-EF slope and dispersal ability.

**Figure 4:**
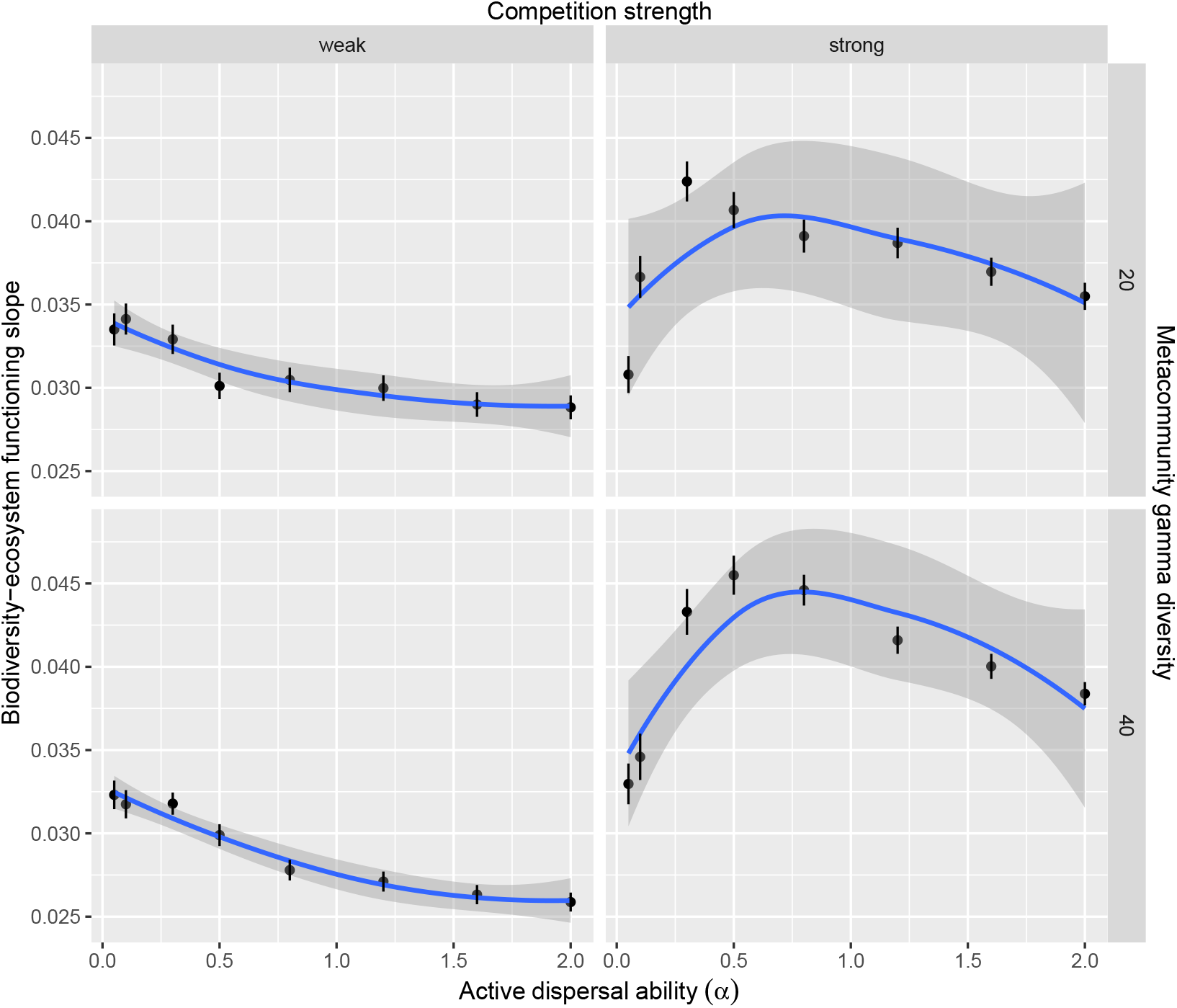
Strength of the biodiversity-EF relationship as a function of dispersal. Columns show weak (left) and strong (right) competition scenarios, while rows show low (top) and high (bottom) metacommunity diversity scenarios. A loess smoother has been added to show the relationships more clearly. When competition is weak, there is a weak relationship between dispersal and biodiversity-EF relationships. In contrast, strong competition facilitates a strong link between biodiversity and EF, with maximum slopes observed at intermediate dispersal rates.

### 3.2 Case study 2: Periphyton community structure in response to changing N:P ratios

For a second case study, we integrate field measurements with an *in silico* experiment to explore how point-source nitrogen pollution could influence biodiversity of periphyton in a network, while using a topology derived from a real river. We use dissolved inorganic nitrogen (DIN, in mg/L) and soluble reactive phosphorous (SRP, mg/L) measured at 31 sites in the Kamp river (Fig. 2). We set the model to separately track N and P concentrations; however, when determining the niche of species, we used the N:P ratio. This ratio can be an important determinant of periphyton community structure in rivers (Stelzer and Lamberti 2001). Moreover, this use highlights the capability of the model to use metrics derived from multiple resources as a single niche dimension.

At the same 31 sites, periphyton community composition was measured by metabarcoding. We scraped off biofilm from the sun-facing side of at least 10 randomly collected stones per site and extracted DNA via DNeasy PowerSoil Pro Kit (Qiagen, Germany). LGC Genomics Gmbh (Berlin, Germany) performed the amplification of the 18S rRNA using primers DIV4for: 5’-GCGGTAATTCCAGCTCCAATAG-3’ and DIV4rev3: 5’-CTCTGACAATGGAATACGAATA-3’ (Visco et al. 2015; Zimmermann et al. 2011; Zimmermann et al. 2015), library preparation (2 x 300 bp) and sequencing on a MiSeq Illumina platform. Finally, species were obtained as amplicon sequence variants (ASVs) using the DADA2 R package (Callahan et al. 2016; Callahan et al. 2017). For complete sampling methods, periphyton sampling methods and water chemistry methods follow (Fuß et al. 2024b) and DNA extraction follows (Fuß et al. 2024a). From the complete community dataset, we retained the 45 most abundant ASVs that were present at at least ten reaches. For modelling, niche optima and breadths were computed as the abundance-weighted mean and sd of the N:P ratio for each taxon.

To establish initial conditions for resources in the network, we used measured values if available, and performed discharge-weighted linear interpolation for all other reaches. Boundary conditions for resources (i.e., the concentrations of resources from lateral input) were identical to the starting conditions. We also assumed that algae dispersed from outside the river network (e.g., via overland flow, animal-mediated dispersal, wind, etc). We computed out-of-network dispersal rates proportionally to the downstream growth in catchment area; upstream reaches with small catchments had low outside dispersal flux, relative to dispersal into downstream reaches.

We conducted this experiment in a similar manner as in case study one, setting up simulations for scenarios (replicated twenty times each) resulting from the variation of two parameters. First, we varied location to see if the topology of the scenario had any impact on how point source pollution affected diversity throughout the network. For pollution sources, we used one reach very near the headwaters (reach 34), and two reaches at intermediate distances from the headwaters but in different subcatchments (reaches 15 and 41; see Fig. 2). Second, we varied the input concentration of N, with levels of 15 and 60 mg/L, equivalent to roughly 0.5 and 2 times the maximum concentrations for wastewater discharge in the European Union (Fernández-Nava et al. 2008; Directive 91/271/EEC 1991). We also ran a control scenario without pollution for comparison.

We ran the model at a daily time scale allowing for 100 days of equilibration at initial conditions. We then applied the pollution treatment and ran the model for an additional 130 days at the new conditions, the first 100 of which were discarded to allow the system to equilibrate. We report results as averages across all twenty replicates and across the final 30 days of simulation. All code for running the scenarios, as well as a copy of the flume package, are archived on Figshare.

All pollution treatments had a strong negative effect on ASV richness at the source, and in most cases these effects persisted downstream, although the strength of the effect decreased with distance from the source (Figs. 5, 6). High pollution scenarios essentially eliminated most taxa from all downstream reaches. Low pollution scenarios similarly eliminated most taxa from the polluted reach and those immediately downstream. However, once an unpolluted tributary entered the polluted reach, richness increased, in many cases exceeding the richness in the control scenario. This suggests there may be a fertilisation effect, whereby the right combination of nutrient input and dispersal can facilitate more diverse communities than what is found under “natural” conditions. Interestingly, in some scenarios, we also observed a slight decrease in richness upstream from the source, suggesting that, with these parameters, continuity of habitat is important for facilitating upstream dispersal (Fig. 6, middle panel).

**Figure 5:**
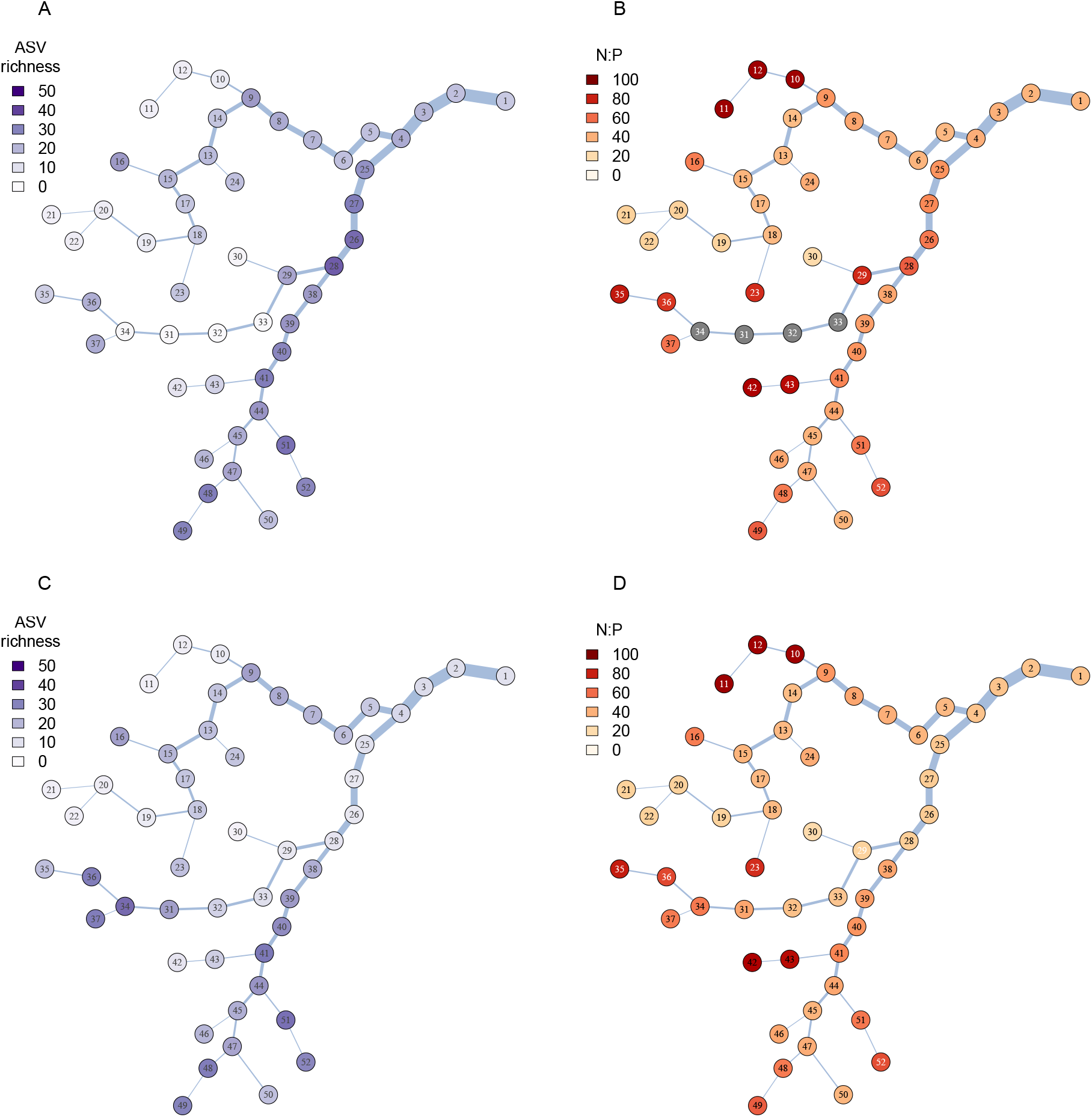
Comparison of periphyton asv richness (A, C) and pollution concentrations (B, D) for the low pollution treatment (A, B; N source concentration = 15 mg/L) and control (C, D). Pollution effects and associated changes in richness persist along the entire length of the stream from the source (reach 34). Reaches with N:P ratios > 100 (i.e., beyond the lethal range for all species) are shown in grey (N:P ratio for these reaches ranged from 147 to 332).

**Figure 6:**
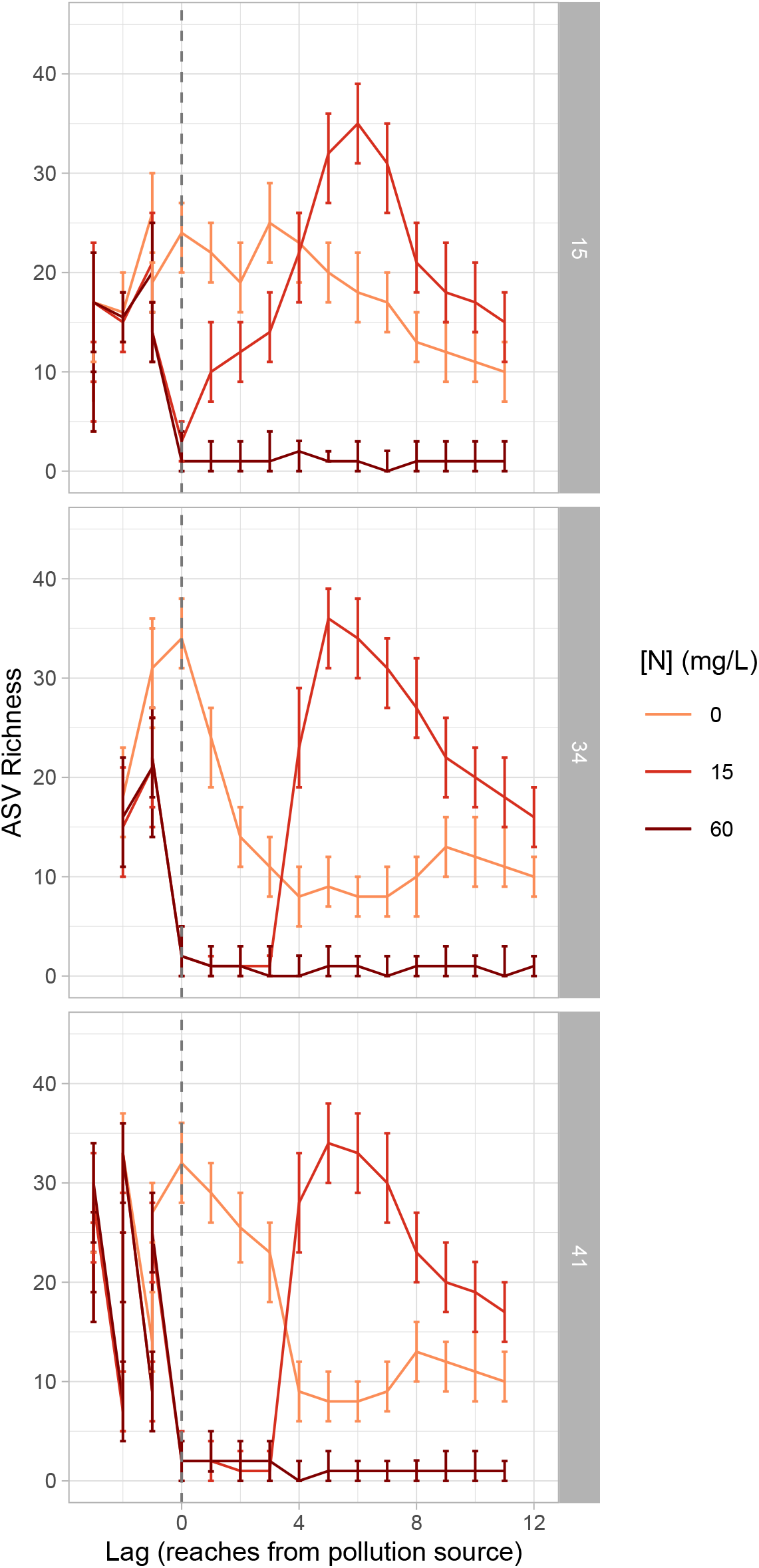
Comparison of asv richness across all scenarios. Lag values indicate the number of reaches downstream of the pollution source (which is marked with a dashed line). Rows indicate the reach where pollution was applied. Error bars show 95% quantile ranges across repeated simulations. Only reaches directly connected to the polluted reach are shown. Negative lag values indicate reaches upstream from the pollution source.

To examine how pollution affected periphyton community composition, we computed Jaccard dissimilarity using a presence-absence matrix. We considered a taxon to be locally present if it occurred in a reach in at least 8 of the last 30 days of the simulation. We took the median Jaccard value by reach pair across replicate simulations for each scenario. For visualisation, we then performed non-metric multidimensional scaling (NMDS) on the resulting Jaccard matrix using the R package vegan (Oksanen et al. 2026), and then plotted the first two axes for the low-pollution and control scenarios. Community composition in polluted reaches showed high magnitude changes relative to the same reaches in unpolluted controls as well as to unaffected reaches within the pollution scenario (Fig. 7). Interestingly, dilution from a single tributary was enough to return the periphyton community to similar compositions as to unpolluted reaches.

**Figure 7:**
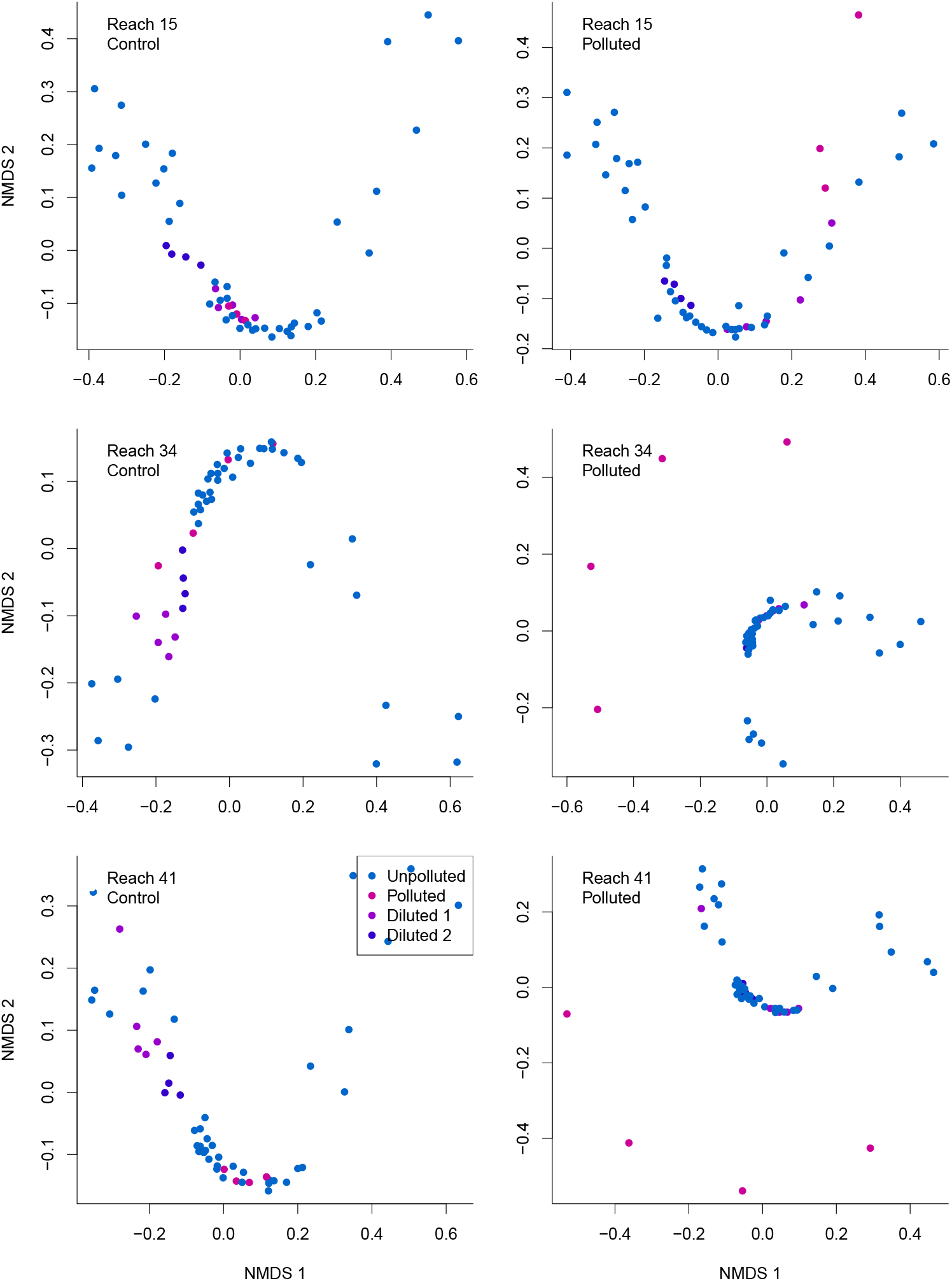
Comparison of community composition between control (left) and pollution (right) treatments for three different pollution origins (rows). In all cases, the weakest pollution treatment (i.e., 15 mg/L) was used. There was little impact on community composition when reach 15 was polluted. For the other two origins, reaches directly affected by pollution unique communities relative to the control, whereas reaches downstream from polluted reaches but diluted by either one or two major tributaries (i.e., “Dilution 1” and “Dilution 2”) were similar to unpolluted reaches.

## 4 Discussion

We have presented a new model for interacting community and resource dynamics in fluvial meta-ecosystems, along with an R-package *flume* implementing the model (Talluto 2023). Our model is based on coupling a patch dynamics model (similar to classical metapopulation models; Levins 1969) with a reaction transport model. The model is incorporates the unique structure of fluvial meta-ecosystems (e.g., local ecosystems connected asymmetrically and mediated by discharge), and includes process-based simulations for borth resource/niche dynamics and the presence-absence of organisms. We demonstrate with our case studies that the model can be used to explore hypotheses about the mechanisms structuring biodiversity-EF relationships and metacommunity structure. In the first case study, the model reproduced a theoretically predicted hump-shaped relationship between dispersal and the biodiversity-EF slope under specific conditions. In particular, the hump shape was only visible when competition was a strong influence in structuring the biological community (Fig. 4). Variation in the output across replicate simulations and due to different species in the metacommunities was low relative to the magnitude of the effects. In the second case study, we partially parametrised the model with data from a field study, then manipulated nutrient inputs to simulate point-source pollution. High levels of pollution had strong effects on both richness and community composition, with effects that reached far beyond the local reach receiving nitrogen input (Figs. 5, **??**, 7). Some partially affected reaches became diversity hotspots, and even non-polluted reaches experienced changes in community composition (Fig. 7).

Movements of resources and organisms in meta-ecosystems can have profound implications for ecosystem structure and function (Tonkin et al. 2018). Nutrient fluxes in particular can mediate diversity and stability in local communities, and can impact the trait distribution within local communities (Gravel et al. 2010). The extent to which ecosystems exchange resources and the degree to which this exchange imposes costs and benefits on each ecosystem can lead to complex dynamics, including alternative stable states (Messan et al. 2018). In rivers, cross-ecosystem fluxes of both organisms and material are facilitated by the flow of water, and are a dominant feature structuring both biological communities and habitats. Thus, meta-ecosystem models in rivers must account for these very high cross-ecosystem flows while also accounting for interactions both within local biological communities and between communities and resources. Moreover, the match of organisms to resource conditions can strongly influence EF (Talluto et al. 2024). Our first case study demonstrates this point, with stronger relationships between biodiversity and EF when high gamma diversity and high dispersal rates facilitate organisms reaching habitats where their fitness is highest (Fig. 4).

### 4.1 Parametrisation

Parametrising the model presents a number of challenges. Although the model parameters describe ecological processes, many of them will be very difficult to measure in practice. For example, measuring dispersal rates is possible, but our model expects dispersal in terms of the contribution to total colonisation flux (and incorporates both the movement of organisms as well as the successful establishment at the colonisation site). This is nearly impossible to measure in the field, although it could be measured or controlled for in laboratory conditions. However, measurement of dispersal in terms of relative rates of upstream and downstream dispersal is possible, as is estimation of dispersal kernels; this could still be useful for calibration by establishing relationships among model parameters.

At a higher level, many of the model’s outputs are frequently measured under field conditions, such as bulk rates of EF or local and regional biodiversity patterns. Thus, inverse calibration becomes possible. Such a procedure would involve obtaining information from field or lab experiments that correspond to particular model outputs (as well as inputs when possible). Then a calibration algorithm can search for parameter combinations that maximise the likelihood of the measured quantities given the model outputs. Because the model has many free parameters, some of which are likely exchangeable, simple optimisation algorithms are unlikely to produce sensible results. However, algorithms that perform better with multiple local optima, such as simulated annealing or Markov Chain Monte Carlo (MCMC) may be fruitful. Bayesian calibration using MCMC in particular has two benefits. First, a modeller can constrain model parameters using prior distributions, helping to narrow the search space and making it less likely that the model produces unrealistic parameter estimates. Second, full posterior inference allows for multiple parameter combinations to be included in the solution set and compared probabilistically. For example, a given data set may reveal that either active dispersal and competition are both very high, or dispersal and competition are very low, but only if there is also immigration from outside the river network. This kind of exercise could prove very useful in suggesting hypotheses and guiding future field studies. A major challenge of this approach is computational; MCMC may require many model runs to fully explore the parameter space, especially for high-dimensional models.

### 4.2 Applications

One of the main applications of our model and the accompanying software is in rapidly exploring hypotheses relating to diversity and EF at the scale of entire river networks. Conducting field or laboratory experiments at the scale of entire networks can be cost-prohibitive, whereas manifold manipulations can easily be performed *in silico* (Tonkin et al. 2018). For example, changes in river network topology (e.g., due to human influence or changing precipitation patterns) can influence the balance between resource consumption and transport, and also change metacommunity dynamics (and associated feedbacks between the community and resource consumption) (Sabo and Hagen 2012; Tonkin et al. 2018). In the field, such changes can be studied either in time (e.g., by studying a single network before and after disruption), or in space (e.g., studying many networks with different spatial properties), but in both cases such studies are expensive and logistically difficult. Our model could be used to explore potentially fruitful avenues of research, to suggest experimental designs, and to evaluate potential mechanisms before committing resources to a field study.

Large-scale flow experiments, often in association with changes in managed flow regimes downstream of large dams, are one way to test connectivity changes across a significant portion of a river network (Konrad et al. 2011). However, these experiments are expensive and necessarily involve multiple stakeholders with potentially competing interests, and not all such experiments will have the resources to collect the relevant ecological data at meta-ecosystem scales (Olden et al. 2014; Kuemmerlen et al. 2019). Moreover, in many cases technical challenges preclude making the appropriate measurements; e.g., sensor limitations or low resolution of monitoring data can result in important changes not being detected (Droujko et al. 2025). Moreover, many experimental manipulations that might be physically feasible (e.g., pollution, changes in land use, invasive species) cannot be performed for ethical reasons due to their environmental impact. Experiments of this sort must be done in the laboratory, where spatial scale is often limiting and scaling rules are poorly understood, or opportunistically, when natural or anthropogenic events produce the desired manipulations. Indeed, when such manipulations are planned (e.g., dam construction or removal), our model can be used to experiment in advance with potential scenarios, informing field data collection or providing information about the range of possible impacts and restoration outcomes.

We have presented a meta-ecosystem model that allows for the coupling of resource dynamics with species distributions. This model allows for in-silico experiments exploring the relationships between metacommunity and biogeochemical processes in FMEs. To demonstrate some of the possible applications of the model, we have presented two case studies, informed by the topology and (for the second case study) community dynamics of a real river. We hope that these examples, along with the relatively user-friendly implementation of the model via an R package, will stimulate further explorations of meta-ecosystem dynamics in FMEs.

## 5 Supplemental Information

**Figure S1:**
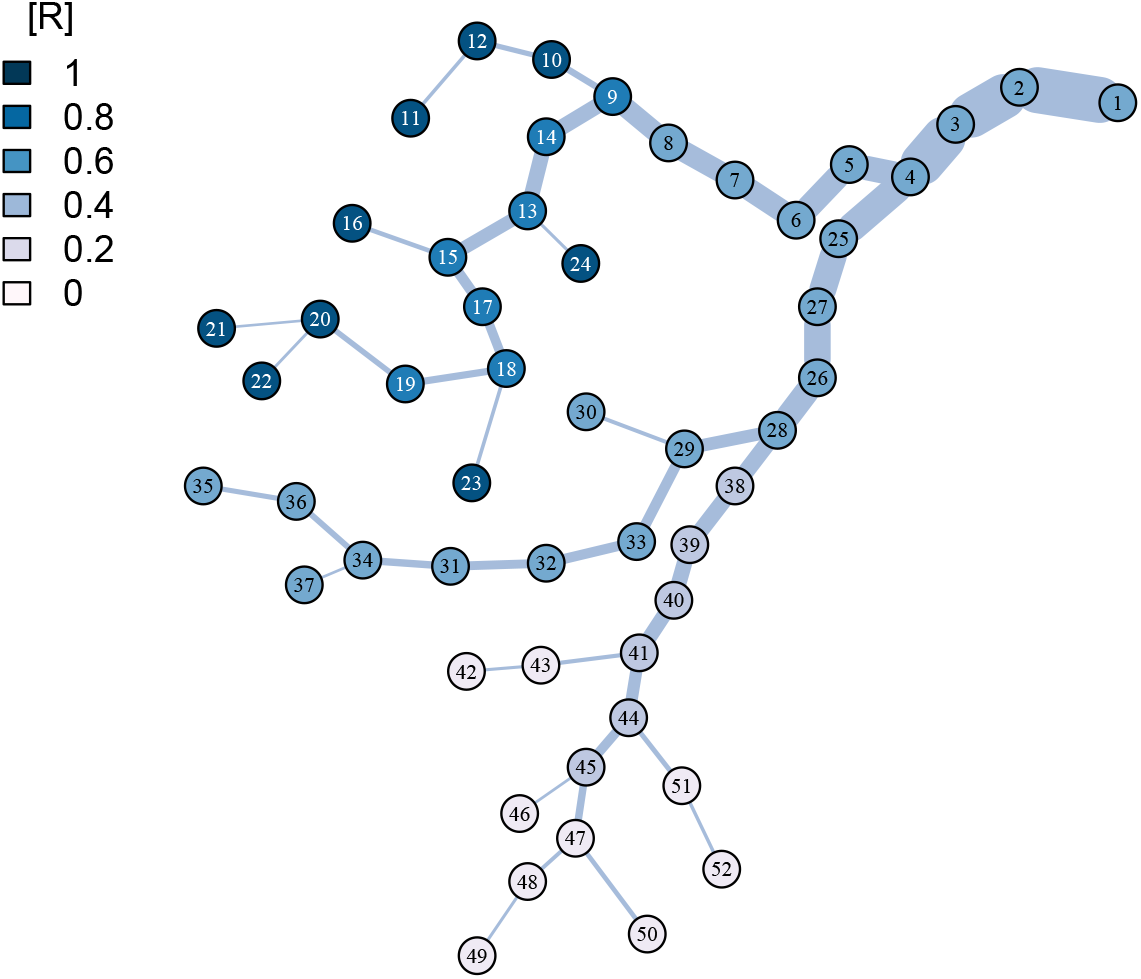
Initial resource state of the river network for the first case study, establishing a resource gradient from high concentrations in the northernmost tributaries to very low concentrations in the southern tributaries. Node colours give the resource concentration ([R]).

